# Immunoglobulin J chain as a non-invasive indicator of pregnancy in the cheetah (*Acinonyx jubatus*)

**DOI:** 10.1101/831628

**Authors:** Michael J. Byron, Diana C. Koester, Katie L. Edwards, Paul E. Mozdziak, Charlotte E. Farin, Adrienne E. Crosier

## Abstract

The North American cheetah population serves as a reservoir for the species, and acts as a research population to help understand the unique biology of the species. Little is known about the intrauterine physiology of the cheetah, including embryo differentiation, implantation, and the development of the placenta. After mating, cheetah females frequently experience (30-65% of matings) a non-pregnant luteal phase where progestogen metabolite levels match those found in pregnant females for the first ~55 days of gestation, but parturition does not occur. Immunoglobulin J chain (IgJ) is a molecule that is involved in the activation of the secretory immune response and has been found to be indicative of pregnancy in the cheetah using fecal monitoring. In this study, western blotting was employed to track IgJ abundance in pooled weekly fecal samples following natural breeding or exogenous stimulation to ovulate, and IgJ levels were compared between individuals undergoing a pregnant (n = 12) and non-pregnant (n = 19) luteal phase. It was revealed that IgJ abundance was increased in pregnant females compared to non-pregnant females at week 4 and week 8 post-breeding, indicating the potential modulation of maternal immunity in response to sensitive events such as implantation and the increased secretory activity of the placenta. IgJ levels also tended to be higher early after breeding in females that were bred naturally with intact males compared to exogenously stimulated females with no exposure to seminal plasma, indicating the promotion of maternal tolerance to seminal antigens present upon embryonic implantation. Monitoring fecal IgJ may be a potential method to determine gestational status in the cheetah and will aid future conservation efforts of the species.

## Introduction

The cheetah (*Acinonyx jubatus*) is listed as vulnerable by the International Union for Conservation of Nature (IUCN), with an estimated population of ~7100 individuals in the wild, and numbers continuously decreasing due to habitat fragmentation and human conflict [1]. Because of the threats to wild cheetahs, an *ex situ* population is critical to serve as an insurance population should the wild cheetah’s numbers diminish further. The *ex situ* population can serve as a reservoir for the species and could potentially be used for reintroduction efforts in the future. The insurance population is also invaluable for research purposes, allowing for studies that cannot be conducted on *in situ* populations due to the scarcity of the species in the wild.

The cheetah is an induced ovulator, meaning that mating or exogenous hormones are necessary for ovulation to occur [2]. While the reproductive events of the domestic cat have been studied in depth [3, 4], little is known about the intrauterine physiology following breeding in wild felids, including the timing of events such as embryo differentiation, implantation, and placentation. Interestingly, cheetahs in human care often encounter reproductive challenges that their wild counterparts do not, as many breedings among cheetahs in human care are unsuccessful. “Pseudopregnancy,” or a non-pregnant luteal phase, has occurred in up to 30% to 60% of matings in North American zoos over recent years (2013-2018; Crosier, personal communication). In these unsuccessful matings, ovulation is confirmed by detectable rises in progestogen metabolites in feces, serum or urine [2, 5]. The concentration of these metabolites is elevated for approximately 55 days, and during this time the hormonal profile of a non-pregnant individual is indistinguishable from a pregnant individual. The high prevalence of a non-pregnant luteal phase after breeding in cheetahs under human care has greatly reduced the reproductive potential of the *ex situ* population and has contributed to the challenge of reaching sustainability due to the impact on the genetic diversity of the population.

Recent advances in mass spectrometry and other proteomic analyses have led to the study of excreted biomarkers as diagnostic or treatment tools in a clinical setting [6]. The production of some biomarkers has been shown to be affected by reproductive events, and certain biomarkers have been found to indicate physiological status such as pregnancy in the domestic dog [7] and several wild canid species [8]. Recently, methods have been developed for the identification of fecal biomarkers of pregnancy in the polar bear [9], and another study in the cheetah identified fecal biomarkers with potential roles in early pregnancy establishment using commercially available antibodies [10]. Koester and colleagues identified a novel biomarker immunoglobulin J chain (IgJ), with increased levels in pregnant individuals and were able to distinguish between pregnant and non-pregnant cheetahs in the 4 weeks following breeding. IgJ is a small polypeptide that serves to regulate polymer formation of Immunoglobulin A (IgA) and Immunoglobulin M (IgM), modulating the secretory activity of these molecules [11, 12]. IgJ functions to provide high levels of avidity to IgA and IgM and facilitates their exocrine transfer to mucosal surfaces [13]. The secretory immunoglobulins that IgJ helps to activate are integral in the response to foreign antigens at surfaces such as the endometrium, and IgJ expression is likely modulated by the unique physiological status of pregnancy. Placental factors that are absent in females undergoing a non-pregnant luteal phase may act to modulate the maternal immune response and affect IgJ abundance, allowing for IgJ monitoring as a method for distinguishing between the gravid and non-gravid states in the cheetah. The objective of the current study was to evaluate the temporal patterns of fecal IgJ abundance over the course of pregnancy in the cheetah to determine the timing of intrauterine events that result in either a successful pregnancy or a non-pregnant luteal phase. Changes in IgJ abundance may indicate maternal immune modulation and could reveal certain events such as implantation and placental development that occur during the establishment of pregnancy early after breeding.

## Materials and methods

### Animals

This study was conducted according to the recommendations in the Guide for the Care and Use of Laboratory Animals of the National Institutes of Health. The female cheetahs included in this study (n=17 individuals) were all housed at accredited Association of Zoos and Aquariums (AZA) institutions within the United States. All subjects were born *ex situ* and managed according to the guidelines developed by the Cheetah Species Survival Plan (SSP). The animals included in the study were female adults from 2 to 12 years of age (mean ± standard error of the mean (SEM) = 6.0 ± 0.6 y). The animals in this study were fed a diet of commercial beef or horse-based meat product (Central Nebraska Packing, Inc., Milliken Meat Products Ltd, or Carnivore Diet 10; Natural Balance Pet Foods Inc., Pacoima, CA) a minimum of five days per week, with supplements that included whole rabbit, beef and horse bone, or organ meat. Water was available *ad libitum*.

Fecal samples were collected opportunistically from females that were naturally bred according to SSP breeding management recommendations and from females receiving exogenous gonadotropins to stimulate ovulation. Exogenous gonadotropin therapy was conducted under IACUC approval for separate projects according to previously published methods [14, 15], and included the stimulation of follicular development (with equine chorionic gonadotropin), followed by stimulation of ovulation (with human chorionic gonadotropin or porcine luteinizing hormone). Pregnancy was confirmed by the birth of offspring, and non-pregnant luteal phase was confirmed by an increase in progestogen metabolite concentration after either natural breeding with no cubs produced or exogenous gonadotropin administration.

### Sample collection and preparation

Fecal samples were collected non-invasively from enclosures approximately 3-4 times weekly. Only fresh samples (deposited within 24 h) were chosen. Samples, about 50g in size, were collected in individual plastic bags and immediately stored in a −20°C freezer. Individual fecal samples were then lyophilized (VirTis, 35L Ultra Super XL-70, Gardiner, NY), crushed, and transferred to individually labeled tubes. Reproductive cyclicity and ovulation were confirmed by steroid hormone metabolite analysis. Fecal samples underwent a steroid hormone metabolite extraction protocol according to previously published methods [16, 10]. Extraction efficiency was determined by the addition of radiolabeled ^3^H-progesterone prior to shaking extraction. The mean extraction efficiency (±SEM) was found to be 73.6% ± 0.2% for all samples.

Weekly pooled fecal samples of 0.5g were created by combining approximately 0.125g of four individual samples in a 15 mL centrifuge tube. Individual samples from day 1-7 post-breeding were used to create the pooled sample for week 1. Individual samples from day 8-14 post-breeding were used to create the pooled sample for week 2, etc. Total protein was subsequently extracted from pooled samples as follows. 6 mL of 0.1 M phosphate buffered saline (0.138 M NaCl, 0.0027 M KCl; pH, 7.4) with protease inhibitor (1:1000) was added to the pooled fecal sample, and the mixture was shaken for 30 min and centrifuged at 4600 x *g* for 30 min. The supernatant was filtered using a 0.22 μm syringe driven filter unit (Millipore Sigma), and the proteins were then precipitated from the supernatant using a 60% ammonium sulfate saturation. The ammonium sulfate solution was shaken for 30 min and centrifuged at 7000 x *g* for 30 min. The protein extract pellet was collected and resuspended in 400 μL of phosphate buffered saline with protease inhibitor. This protein extract solution was then desalted using a 3 kDa Millipore spin column (Amicon Ultra-0.5) and centrifugation at 7400 x *g*. All extraction steps were performed at 4°C. Extracted samples were then run on a Bradford assay (Bio-Rad Protein Assay, Hercules, CA) to determine total protein concentration. Briefly, standards for the assay were created by serial dilution at 0.388 mg/mL to 0.012 mg/mL. Fecal protein samples were diluted to 1:30, and 10 μL of each sample was added to a well. 200 μL of Bio-Rad Quick Start™ Bradford Dye Reagent was added to each well, and after 5 minutes protein concentrations were determined using a Molecular Devices Filtermax F5 plate reader. Differences in steroid hormone and total protein concentrations between pregnant and non-pregnant groups were determined using a Student’s T-test in R (version 3.3.2) [17], with differences considered significant at P < 0.05.

### Steroid hormone metabolite analysis

Steroid hormone neat extracts were diluted 1:20 to 1:16,000 in phosphate buffer (2.2 M NaH_2_PO_4_, 3.5 M Na_2_HPO_4_, 0.3 M NaCl, H_2_O; pH, 7.0) and were run for analysis on enzyme immunoassay (EIA). Estrogen metabolites in diluted fecal extracts were used to determine reproductive cyclicity using an estradiol EIA that has been validated for use in the cheetah [18]. Briefly, for samples collected before 2015, a polyclonal anti-estradiol antibody (R4972; C. Munro, University of California, Davis, CA) was added to a 96-well microtiter plate and incubated for 12 h. Diluted samples, standards, and peroxidase-enzyme conjugated 17β-estradiol were added, and the plate was incubated for 2 h at 23°C. Unbound components were washed off, and an ABTS chromogen solution (2,2’-azino-bis(3-ethylbenzothiazoline-6-sulphonic acid)) was added as a substrate. Optical densities of each well on the plate were determined using a microplate reader (Molecular Devices Filtermax F5, reading filter 405 nm, reference filter 540 nm). Estrogen metabolite concentrations for samples collected from 2015-2018 were determined using a revised protocol for the estradiol EIA [18]. Briefly, diluted samples, standards, peroxidase enzyme conjugated 17β-estradiol, and antibody (R4972; C. Munro, University of California, Davis, CA) were added to a pre-coated goat-anti-rabbit IgG plate, and the plate was incubated for 2 h at 23°C. Unbound components were washed off, a TMB chromogen solution (3,3’, 5,5;-tetramethylbenzidine) was added as a substrate, and the reaction was halted with addition of 1N HCL. Optical densities of each well on the plate were determined using a microplate reader (Molecular Devices Filtermax F5, reading filter 405 nm, reference filter 540 nm). Inter-assay variation was monitored through the use of two internal controls, and coefficients of variation for all samples in duplicate were <10%.

Progesterone metabolites were used to determine ovulation and the presence of a luteal phase. Concentrations were determined using a progesterone EIA that has been validated for use in the cheetah [18], using a monoclonal progesterone antibody (no. CL425, Quidel Co., San Diego, CA), and an associated peroxidase-enzyme conjugated to progesterone (C. Munro, University of California, Davis, CA). Plates were prepared and run using the same procedure as the estradiol assay, with samples run using the revised protocol and pre-coated goat-anti-mouse IgG plates beginning in 2012. Internal controls were used to monitor inter-assay variation, and coefficients for samples in duplicate were <10%. In both cases, fecal hormone metabolite concentrations and profiles were comparable across the two EIA protocols.

### Western blotting and protein quantification

Total protein samples were diluted to 2 mg/mL in MilliQ water to a final volume of 30 μL. Human recombinant IgJ (Abcam #140727) was used as a positive control at 16.67 μg/mL. Samples were then separated by SDS-PAGE, transferred to a PVDF membrane, blocked with 5% milk, and incubated overnight at 4°C with a primary antibody (Aviva Systems Biology ARP55440_P050) diluted 1:1000 in 1% milk. This polyclonal antibody was developed in a rabbit against human recombinant IgJ and was previously found to be reactive to cheetah IgJ in western blot [10]. The membrane was then incubated with a secondary antibody (Cell Signaling Technology, Anti-Rabbit IgG HRP-linked antibody, #7074S) diluted 1:2500 in 1% milk, and then incubated with a chemiluminescent substrate (Bio-Rad, Clarity Max Western ECL Substrate, #1705062). Membranes were imaged on a G:Box Chemi XRQ (Syngene). Coomassie staining and image analysis of total protein were conducted in order to serve as a loading control [10, 19].

Intensity of IgJ abundance was determined using GeneSys Spot Blot analysis of the band occurring within each lane at 18 kDa for each weekly pooled sample. GeneSys Total Lane analysis for each sample of the Coomassie image was used to determine the total protein in the sample as a loading control. A ratio of IgJ intensity to Coomassie intensity was calculated for each pooled sample, as well as for the positive control. Relative intensity for each pooled sample was calculated by dividing the ratio for each pooled sample by the ratio of the positive control, in order to control for inter-blot variation. A pre-breeding relative intensity value specific to each individual luteal phase was then subtracted from each weekly post-breeding relative intensity value to control for pre-breeding IgJ levels. If relative intensity values were not normally distributed, then the data was subjected to a log transformation. After transformation, 15 of 18 groups were verified for normality using a Shapiro-Wilk test in R (version 3.3.2) [17]. Differences in IgJ intensity between pregnant and non-pregnant groups were determined using Student’s T-test or Mann-Whitney U-Test in R (version 3.3.2) depending on normality of the group, with differences considered significant at P < 0.05, and differences considered a tendency at 0.05 ≤ P < 0.09.

## Results

### Fecal steroid metabolite and total protein concentrations

Fecal estrogen metabolite profiles confirmed the cyclicity of monitored females. Examples of the estrogen metabolite concentrations of cycling females prior to natural breeding can be seen in Fig 1a and 1b. Fecal progestogen metabolite concentrations were significantly higher (P < 0.01) during pregnancy and non-pregnant luteal phase than during a pre-breeding period (Table 1), confirming the presence of a luteal phase after breeding. Fecal progestogen profiles of pregnant females can be distinguished from non-pregnant luteal phase females after around 55 days post-breeding, when progestogen concentrations of non-pregnant females drop (Fig 1c), while pregnant females have extended progestogen excretion until parturition (Fig 1d). Mean total protein concentration of sample extracts from pregnancies (5.93 ± 0.29 mg/mL), non-pregnant luteal phases (6.24 ± 0.23 mg/mL), and pre-breeding (5.69 ± 0.57 mg/mL) were not significantly different (P > 0.05).

**Table 1.**
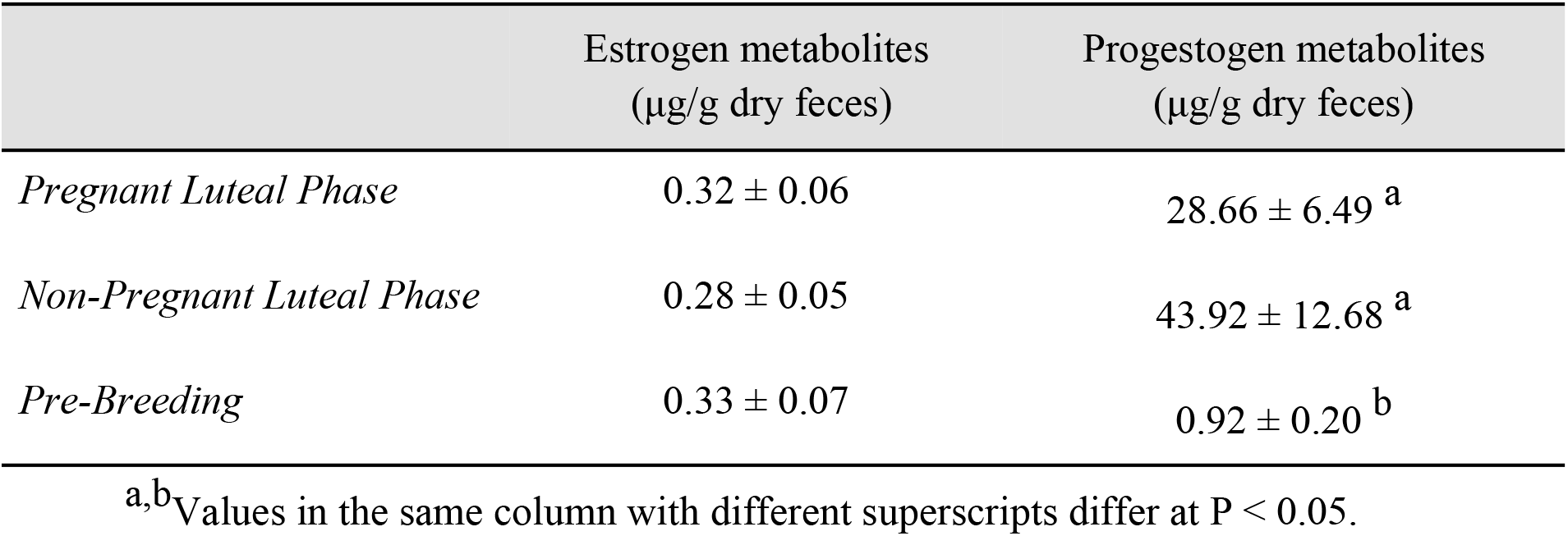
Mean (±SEM) cheetah estrogen and progestogen fecal metabolite concentrations.

**Fig 1.**
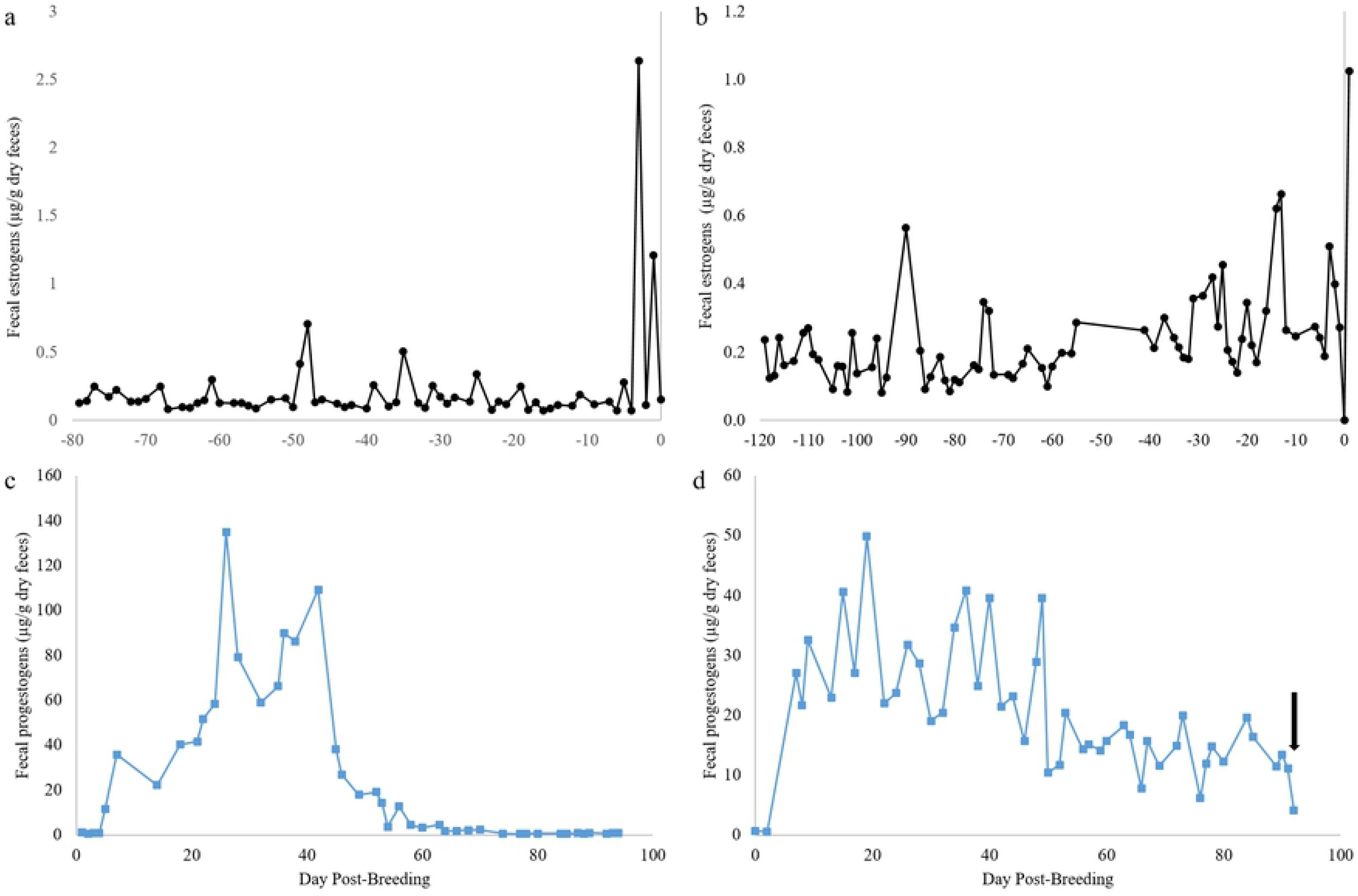
Cheetah estrogen and progestogen fecal metabolite profiles. (a) Estrogen fecal metabolite profile of a cycling female prior to natural breeding and successful pregnancy. (b) Estrogen fecal metabolite profile of a cycling female prior to natural breeding and non-pregnant luteal phase (Axes are different scales). (c) Progestogen fecal metabolite profile of a non-pregnant luteal phase after natural breeding. Progestogen metabolite concentrations return to pre-ovulatory levels around ~55 days post-breeding. (d) Progestogen fecal metabolite profile of a pregnant female after natural breeding. Progestogen fecal metabolite concentrations remain elevated until parturition (Parturition indicated by arrow) (Axes are different scales).

### Post-breeding IgJ response in females exposed to seminal plasma during natural breeding

Detection of IgJ by western blotting with the use of a commercially available antibody was confirmed by the use of a positive control. IgJ was confirmed in the positive control and in fecal samples at a molecular weight of ~18 kD (Fig 2). Females that were bred naturally by intact males with semen deposition, including both successful pregnancies and non-pregnant luteal phases, tended to have significantly higher peak IgJ levels (P = 0.076, mean ± SEM = 0.86±0.05) in the 2 weeks immediately following breeding compared to exogenously stimulated females that did not have exposure to seminal plasma (0.75±0.02) (Fig 3). An example of a post-breeding response can be seen in Fig 2a, with high IgJ abundance in week 1 and week 2 following the female’s first breeding and exposure to seminal plasma. A response from a second female can be seen in Fig 2b, with high IgJ abundance in week 2 following natural breeding and exposure to seminal plasma. In females that were exogenously stimulated to ovulate without exposure to seminal plasma, no increase in IgJ was seen in the two weeks following breeding (Fig 2c).

**Figure 2.**
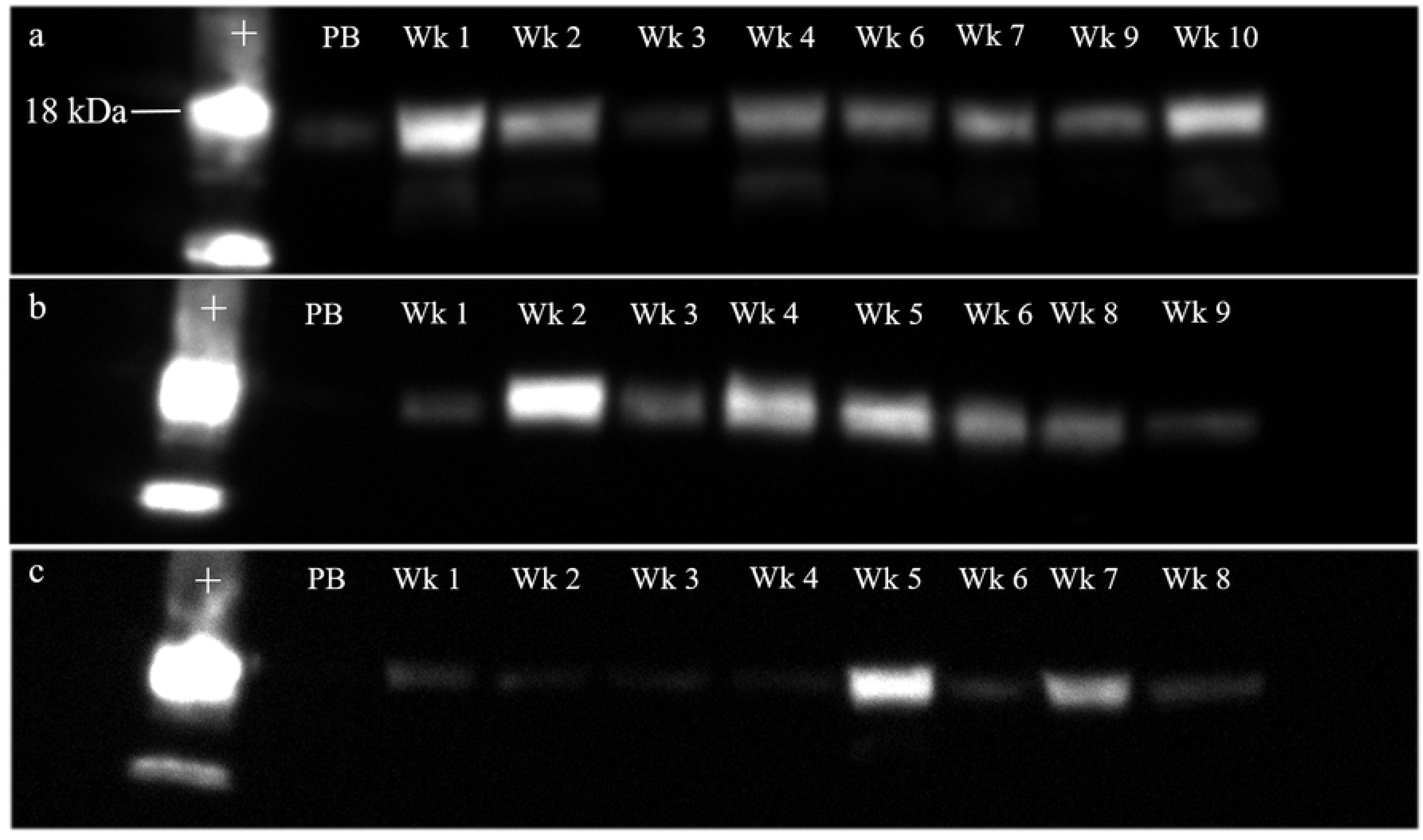
IgJ response immediately following breeding in the cheetah. (a) A female (#6593) that was bred naturally with exposure to seminal plasma had high IgJ levels in week 1 and week 2 post-breeding. (b) A female (#6462) that was bred naturally with exposure to seminal plasma had high IgJ levels in week 2 post-breeding. (c) A female (#8957) with no exposure to seminal plasma following exogenous stimulation to ovulate did not experience an increase in IgJ levels in weeks 1 or 2 following breeding. Positive control is denoted by “+.” Pre-breeding sample is denoted by “PB.”

**Fig 3.**
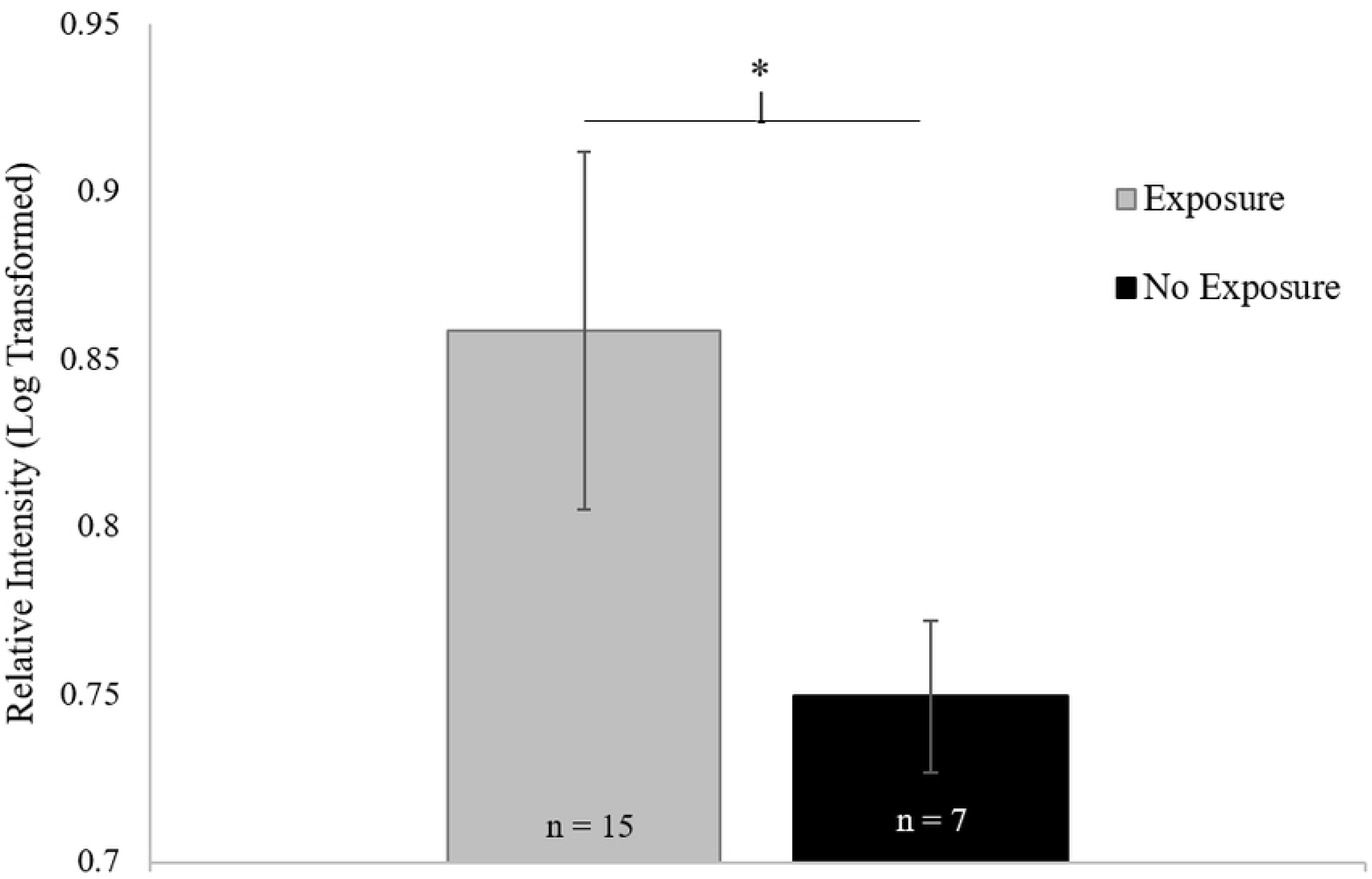
Effect of exposure to seminal plasma on IgJ abundance in cheetahs. Females that were bred naturally by intact males and exposed to seminal plasma (n = 15) tended to have higher IgJ levels in the two weeks following breeding compared to females exogenously stimulated to ovulate with no exposure to seminal plasma (n = 7, P = 0.076).

### Temporal tracking of IgJ abundance

Pregnant females tended to have higher IgJ levels (P = 0.081) in week 4 post-breeding (0.83 ± 0.07, n=10) compared to females experiencing a non-pregnant luteal phase (0.69 ± 0.02, n=18) (Fig 4). Pregnant females were also found to have significantly higher IgJ levels (P < 0.05) in week 8 post-breeding (0.79 ± 0.03, n=9) compared to females experiencing a non-pregnant luteal phase (0.69 ± 0.03, n=15). IgJ abundance was not different between the two groups in weeks 1, 2, 3, 5, 6, 7, or 9 (P > 0.1) (Fig 4). An example of a pregnancy can be seen in Fig 5a, with high IgJ levels in Week 4 and following Week 7 post-breeding. An example of a non-pregnant luteal phase can be seen in Fig 5b, with low IgJ levels at or near pre-breeding levels throughout the sample period.

**Fig 4.**
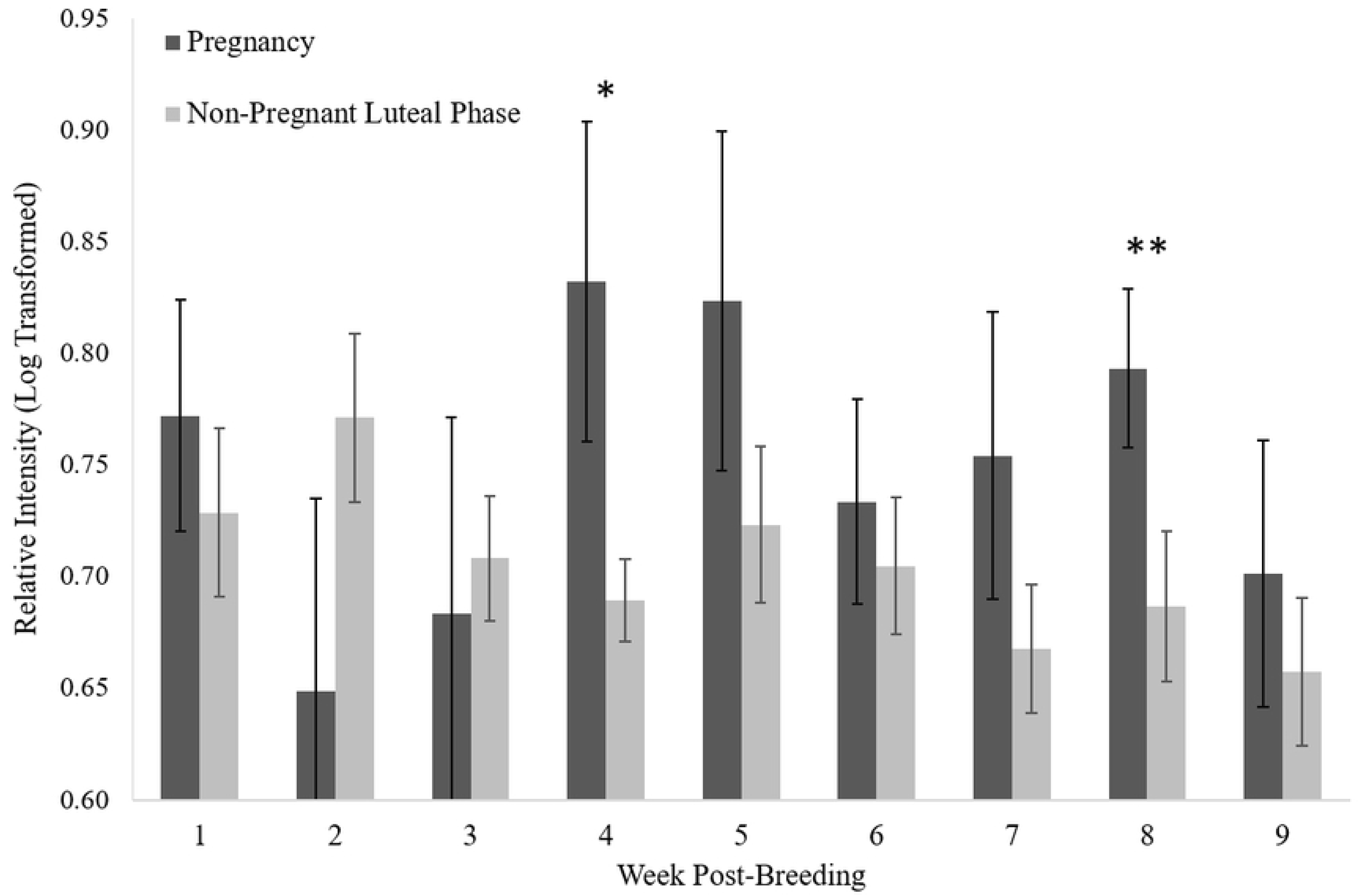
Mean (± SEM) relative intensity of IgJ following breeding in the cheetah. IgJ abundance tended to be higher (P = 0.081 *) in pregnant females (n = 12) compared to those experiencing a non-pregnant luteal phase (n = 19) at week 4 post-breeding. IgJ abundance was significantly higher (P < 0.05 **) for pregnant females compared to during a non-pregnant luteal phase at week 8 post-breeding.

**Fig 5.**
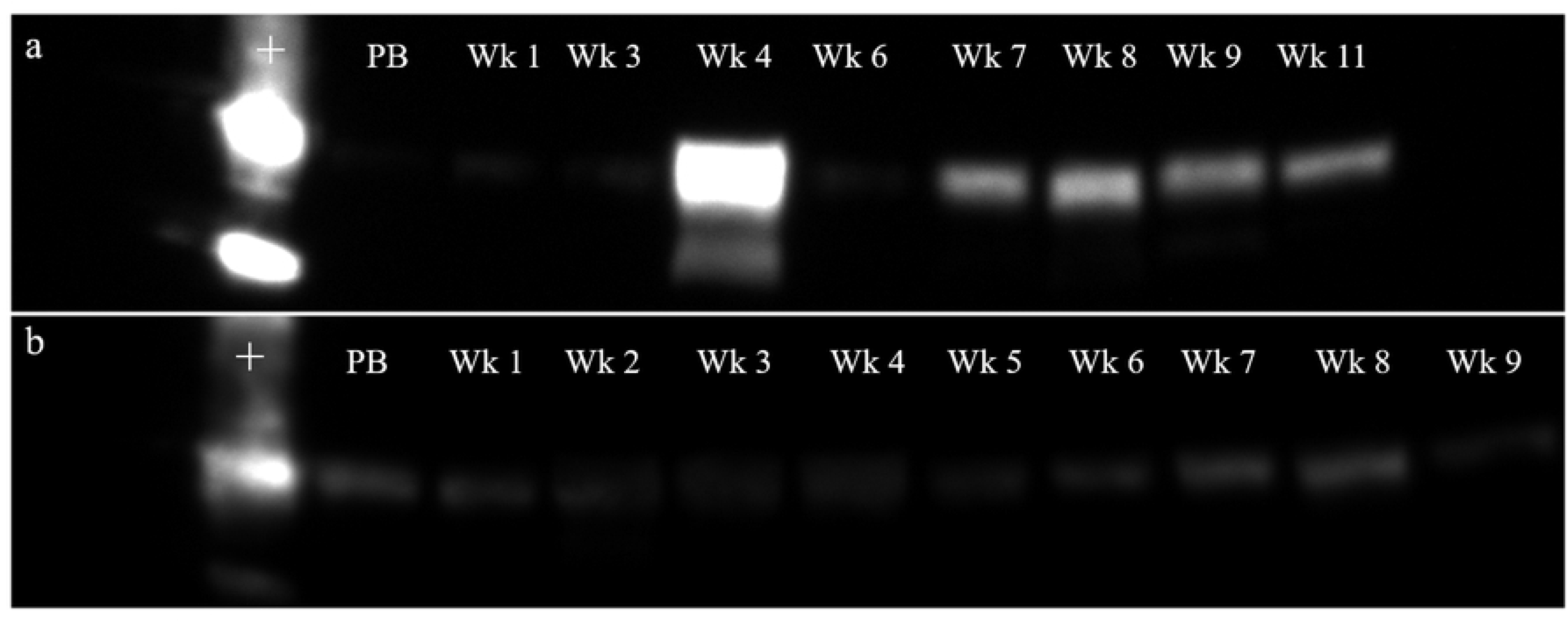
Comparison of pregnant and non-pregnant luteal phase in the cheetah. (a) IgJ was increased in week 4 and following week 7 post-breeding during pregnancy (Female #6593). (b) IgJ remained at or near pre-breeding levels throughout the 9 week sample period during a non-pregnant luteal phase (Female #6339). Positive Control is denoted by “+.” Pre-breeding sample is denoted by “PB.”

## Discussion

The reproductive biology of the cheetah has been studied for decades, with great advances in the understanding of this species both *in situ* and *ex situ*. However, cheetahs in the wild are in decline, as habitat fragmentation and human conflict have reduced the natural range of the species [1, 20]. Because of the vulnerable status of the cheetah, it has become an important goal of conservationists to create an *ex situ* insurance population. However, creating a sustainable *ex situ* population with the goal of improving genetic diversity and ensuring future health and adaptability of the species has become a challenge, as many cheetahs struggle to successfully reproduce in human care. The low genetic diversity of the species as a whole, and high levels of inbreeding depression, have contributed to many health and reproductive issues that affect the cheetah, including the impairment of genes mediating immune defenses [21], low fecundity, and the poor semen quality of males both in captivity and in the wild [22, 23]. While wild cheetahs appear to face similar obstacles in terms of inbreeding depression and low genetic diversity, they are able to reproduce with greater success than cheetahs under human care [24, 25], indicating the possibility that *ex situ* environmental factors may be having a negative effect on reproductive capacity. Of importance to animal managers is understanding the relatively high incidence of non-pregnant luteal phase that occurs in the *ex situ* population, whether this represents failure to conceive or early embryo loss.

Temporal tracking of IgJ abundance over the first 9 weeks post-breeding provides insight into the intrauterine events that occur after the success or failure to establish pregnancy in the cheetah. Females in our study demonstrated an elevation of IgJ after natural breeding with an intact male and seminal exposure compared to exogenous stimulation for ovulation with no seminal exposure. One explanation is that a secretory immune response was stimulated by the presence of seminal plasma in the reproductive tract which, as a foreign substance interacting with a mucosal surface, could induce an upregulation in IgJ. An immune response to semen has been documented previously in mice, as lymphocyte synthesis and cytokine activation is triggered in response to the constituents of seminal plasma [26]. The immune response to semen may help to promote active maternal tolerance of paternal antigens of the fetus at the implantation site [26, 27], preventing rejection of the fetus. Upon successful breeding and exposure to semen, it is possible that the females in this study were experiencing a subsequent activation of the secretory immune response in order to promote tolerance of paternal antigens upon implantation of the fetus and invasion of the endometrium. In contrast to naturally bred females, all eight females that were exogenously stimulated to ovulate demonstrated IgJ levels that were at or near pre-breeding levels in the first two weeks post-breeding, indicating that the IgJ response is not due to ovulation, and that natural breeding and successful deposition of semen are needed to see this immune response.

The timing of early intrauterine events, including implantation and the development of the placenta, is unknown in wild felids. In the domestic cat, fertilization takes place in the oviduct up to 48 h after ovulation, and implantation occurs at day 13-14 post-breeding [3]. It is possible that implantation in the cheetah occurs at a different time as the domestic cat (day 13-14 of ~65 d gestation), at a proportional point in the longer ~93 day gestation of the cheetah (day 19-21). In the present study, IgJ levels tended to be elevated in pregnant cheetahs compared to non-pregnant cheetahs during the pooled fourth week after breeding (day 22-28). Increased IgJ levels during the fourth week post-breeding could indicate an activated secretory immune response to the invasion of the endometrium by the embryo following implantation. IgJ regulates the polymer formation of IgA and is a crucial component of the secretory form, facilitating the exocrine secretion of the molecule. Increased IgA secretion likely alters dendritic cell function, leading to the promotion of regulatory T cell expansion and inhibiting the release of inflammatory cytokines, preventing an inflammatory response to paternal antigens present on the fetal trophoblast [28]. These findings suggest that implantation could occur directly before week 4 post-breeding (day 19-21), as a sustained secretory immune response at the endometrium likely continues and intensifies as the implanted embryo continues to develop during the fourth week.

Pregnancy in the cheetah first becomes distinguishable from a non-pregnant luteal phase using progestogen metabolite monitoring at ~day 55 post-breeding, as progestogen levels in fecal samples drop to pre-ovulatory levels in non-pregnant individuals. It can be theorized that around this time in non-pregnant individuals a regression of the corpora lutea (CL) occurs which results in a subsequent decrease in progesterone production, followed by a plateau at lower basal levels. It is unknown whether luteal regression occurs in pregnant females as well, or if the CL is maintained until parturition. Recent studies have suggested that the placenta is a source of progesterone production in the domestic cat [29, 30], and acts to supplement luteal progesterone later in gestation. A placental source of progesterone secretion is likely in the cheetah and may explain the observed increase in IgJ levels at week 8 (day 50-56) post-breeding. An increased and sustained secretion of progesterone from the fetal-maternal barrier at day 50-56 in gestation may modulate secretory immunity, causing IgJ levels to be increased in pregnant individuals compared to non-pregnant individuals. Local placental synthesis of progesterone may act as an immunosuppressive factor at the site of embryonic implantation during both murine and human gestation [31]. Progesterone suppresses T lymphocyte proliferation and inhibits natural killer cell activity at the maternal-fetal interface [32], decreasing the activity of these immune molecules to protect the semi-allogeneic fetus and preventing an inflammatory immune reaction. Because the endometrium is a mucosal surface, it is likely that secretory immunity is modulated as well, promoting tolerance of the fetus and allowing for non-inflammatory neutralization of foreign pathogens. Secretory immunity may also be modulated by the secretion of Prostaglandin F_2ɑ_ (PGF_2ɑ_) by the placenta during week 8 post-breeding, as levels of a fecal PGF_2ɑ_ metabolite (PGFM) were found to be elevated in the pregnant cheetah beginning at day 48 post-breeding and increased through parturition [33]. While PGF_2ɑ_ is known to have a strong luteolytic effect in many species [34], suggesting a possible impact on immunity, the action of this molecule is unknown in felids and the impact on IgJ levels has not yet been determined.

It is possible that the presence of external immune stressors, which potentially could alter the production of systemic IgJ, could influence the results and interpretation of our experiment. IgJ is a protein that is upregulated in response to an activation of the secretory immune system, which is present in all mucosal surfaces of the body in order to protect from foreign pathogens. An immune challenge to a mucosal surface that is independent of pregnancy could increase expression of IgJ, resulting in high IgJ values in our assay that do not correspond to intrauterine events and could affect the accuracy of the assay. Inter-individual variability in IgJ expression was controlled for here by assessing relative changes in concentration from pre-breeding levels within each individual. A future goal is to extend the Western analysis to a reliable benchtop enzyme-linked immunosorbent assay (ELISA) for measuring fecal IgJ levels after breeding. Daily quantification using this method would improve the accuracy and efficiency of IgJ monitoring and may be able to reveal moments of immune challenge that impact establishment of pregnancy with greater precision. The occurrence of a non-pregnant luteal phase is common in other species, indicating the potential for a felid-wide or carnivore-wide assay for determining pregnancy using IgJ monitoring.

In summary, this study detailed IgJ levels throughout the first 9 weeks of pregnancy and non-pregnant luteal phase in the cheetah. Females that were bred naturally and exposed to seminal plasma tended to have a higher immediate IgJ response than females that were exogenously stimulated to ovulate with no seminal exposure, indicating an immune response to the constituents of seminal plasma that may have the effect of promoting maternal tolerance of fetal tissue upon implantation. There was a tendency towards increased IgJ abundance at week 4 post-breeding in pregnant cheetahs compared to non-pregnant cheetahs, indicating an activation of the secretory immune system in response to implantation and the invasion of the maternal endometrium by the fetal trophoblast. A significant increase in IgJ abundance was also found in week 8 post-breeding in pregnant cheetahs compared to non-pregnant cheetahs. Taken together, these data support the suggestion that the window of implantation in the cheetah is between 19-21 days post-breeding, and that the placenta is a source of extragonadal progesterone during the third trimester. These findings will help to improve *ex situ* management of the species, and further research will continue to advance the understanding of cheetah reproductive physiology following breeding, aiding future conservation efforts for the species.

## Acknowledgments

This study was conducted under a consortium agreement of the Conservation Centers for Species Survival (C2S2), a formal partnership that shares unique resources to improve the biological understanding and management of endangered species, especially those that require space, natural group sizes, and scientific research. The authors would like to thank the Smithsonian National Zoological Park, Fossil Rim Wildlife Center, Wildlife Safari, White Oak Conservation, The Wilds, Cincinnati Zoo, Birmingham Zoo, Denver Zoo, Columbus Zoo, Caldwell Zoo, and Omaha Zoo for participation in this study. We would also like to thank Amber Dedrick, Adri Kopp, Jaelyn Currant, and Kelly Mickael for technical support.

## References

1. Durant SM, Mitchell N, Groom R, Pettorelli N, Ipavec A, Jacobson AP, et al. The global decline of cheetah *Acinonyx jubatus* and what it means for conservation. PNAS. 2017;114(3): 528–533.

2. Brown JL, Wildt DE, Wielebnowski N, Goodrowe KL, Graham LH, Wells S, et al. Reproductive activity in captive female cheetahs (*Acinonyx jubatus*) assessed by faecal steroids. Journal of Reproduction and Fertility. 1996;106(2): 337–346.

3. Denker HW, Eng LA, Hamner CE. Studies on the early development and implantation in the cat. II. Implantation: proteinases. Anatomy and embryology. 1978;154(1): 39–54.

4. Leiser R, Koob B. Development and characteristics of placentation in a carnivore, the domestic cat. The Journal of experimental zoology. 1993;266(6): 642–656.

5. Wildt DE, Brown JL, Bush M, Barone MA, Cooper KA, Grisham J, et al. Reproductive status of cheetahs (*Acinonyx jubatus*) in North American Zoos: The benefits of physiological surveys for strategic planning. Zoo Biology. 1993;12(1): 45–80.

6. Burke HB. Predicting clinical outcomes using molecular biomarkers. Biomarkers in Cancer. 2016;8.

7. Kuribayashi T, Shimizu M, Shimada T, Honjyo T, Yamamoto Y, Kuba K, et al. Alpha 1-acid glycoprotein (AAG) levels in healthy and pregnant beagle dogs. Experimental animals. 2003;52(5): 377.

8. Bauman JE, Clifford DL, Asa CS. Pregnancy diagnosis in wild canids using a commercially available relaxin assay. Zoo Biology. 2008;27(5): 406–413.

9. Curry E, Stoops MA, Roth TL. Non-invasive detection of candidate pregnancy protein biomarkers in the feces of captive polar bears (*Ursus maritimus*). Theriogenology. 2012;78(2): 308–314.

10. Koester DC, Wildt DE, Maly M, Comizzoli P, Crosier AE. Non-invasive identification of protein biomarkers for early pregnancy diagnosis in the cheetah (*Acinonyx jubatus*). PLoS One. 2017;12(12): e0188575.

11. Brandtzaeg P. Immunohistochemical characterization of intracellular J-chain and binding site for secretory component (SC) in human immunoglobulin (Ig)-producing cells. Molecular Immunology. 1983;20(9): 941–966.

12. Halpern MS, Koshland ME. Noval subunit in secretory IgA. Nature. 1970;228(5278): 1276.

13. Johansen F, Braathen R, Brandtzaeg P. The J chain is essential for polymeric Ig receptor-mediated epithelial transport of IgA. The Journal of Immunology. 2001;167(9): 5185–5192.

14. Howard JG, Roth TL, Byers AP, Swanson WF, Wildt DE. Sensitivity to exogenous gonadotropins for ovulation induction and laparoscopic artificial insemination in the cheetah and clouded leopard. Biology of Reproduction. 1997;56(4): 1059.

15. Pelican KM, Wildt DE, Pukazhenthi B, Howard J. Ovarian control for assisted reproduction in the domestic cat and wild felids. Theriogenology. 2006;66(1): 37–48.

16. Brown JL, Wasser SK, Wildt DE, Graham LH. Comparative aspects of steroid hormone metabolism and ovarian activity in felids, measured noninvasively in feces. Biology of Reproduction. 1994;51(4): 776–786.

17. R Core Team. R: A language and environment for statistical computing. R Foundation for Statistical Computing, Vienna, Austria. 2016. URL https://www.R-project.org/.

18. Crosier AE, Comizzoli P, Baker T, Davidson A, Munson L, Howard J, et al. Increasing age influences uterine integrity, but not ovarian function or oocyte quality, in the cheetah (*Acinonyx jubatus*). Biology of reproduction. 2011;85(2): 243.

19. Welinder C, Ekblad L. Coomassie Staining as Loading Control in Western Blot Analysis. Journal of Proteome Research. 2011;10(3): 1416–1419.

20. Marker LL, Dickman AJ, Jeo RM, Mills MGL, Macdonald DW. Demography of the Namibian cheetah, *Acinonyx jubatus jubatus*. Biological Conservation. 2003;114(3): 413–425.

21. O’Brien SJ, Johnson WE, Driscoll CA, Dobrynin P, Marker L. Conservation genetics of the cheetah: lessons learned and new opportunities. The Journal of heredity. 2017;108(6): 671–677.

22. Wildt DE, O’Brien SJ, Howard JG, Caro TM, Roelke ME, Brown JL, et al. Similarity in ejaculate-endocrine characteristics in captive versus free-ranging cheetahs of two subspecies. Biology of Reproduction. 1987;36(2): 351.

23. Crosier AE, Marker L, Howard J, Pukazhenthi BS, Henghali JN, Wildt DE. Ejaculate traits in the Namibian cheetah (*Acinonyx jubatus*): influence of age, season and captivity. Reproduction, Fertility and Development. 2007,;19(2): 370–382.

24. Laurenson MK, Caro TM, Borner M. Female cheetah reproduction. National Geographic Research & Exploration. 1992;8(1): 64–75.

25. Kelly MJ, Laurenson MK, FitzGibbon CD, Collins DA, Durant SM, Frame GW, et al. Demography of the Serengeti cheetah (*Acinonyx jubatus*) population: the first 25 years. Journal of Zoology. 1998;244(4): 473–488.

26. Johansson M, Bromfield JJ, Jasper MJ, Robertson SA. Semen activates the female immune response during early pregnancy in mice. Immunology. 2004;112(2): 290–300.

27. Robertson SA, Sharkey DJ. The role of semen in induction of maternal immune tolerance to pregnancy. Seminars in Immunology. 2001;13(4): 243–254.

28. Monteiro RC. Immunoglobulin A as an anti-inflammatory agent. Clinical & Experimental Immunology. 2014;178: 108–110.

29. Tsutsui T, Suzuki Y, Toyonaga M, Oba H, Mizutani T, Hori T. The role of the ovary for the maintenance of pregnancy in cats. Reproduction in domestic animals. 2009;44 Suppl 2(s2): 120–124.

30. Siemieniuch MJ, Jursza E, Szostek AZ, Skarzynski DJ, Boos A, Kowalewski MP. Steroidogenic capacity of the placenta as a supplemental source of progesterone during pregnancy in domestic cats. Reproductive biology and endocrinology. 2012;10(1): 89.

31. Siiteri PK, Stites DP. Immunologic and endocrine interrelationships in pregnancy. Biology of Reproduction. 1982;26(1): 1–14.

32. Szekeres-Bartho J. Immunological relationship between the mother and the fetus. International Reviews of Immunology. 2002;21(6): 471–495.

33. Dehnhard M, Finkenwirth C, Crosier A, Penfold L, Ringleb J, Jewgenow K. Using PGFM (13,14-dihydro-15-keto-prostaglandin F2α) as a non-invasive pregnancy marker for felids. Theriogenology. 2012;77(6): 1088–1099.

34. Senger PL. Pathways to pregnancy and parturition. 2nd rev. ed. Pullmann, Wash: Current Conceptions; 2005.

